# Botulinum toxin intoxication requires retrograde transport and membrane translocation at the ER in RenVM neurons

**DOI:** 10.1101/2023.10.17.562749

**Authors:** Jeremy C. Yeo, Felicia P. Tay, Rebecca Bennion, Omar Loss, Jacquie Maignel, Laurent Pons, Keith Foster, Matthew Beard, Frederic Bard

**Affiliations:** Institute of Molecular and Cell Biology, 61 Biopolis Drive, Singapore 138673; Ipsen Bioinnovation, 102 Park Drive, Milton Park, Abingdon OX14 4RY, UK; Ipsen Innovation, 5 Avenue du Canada, 91940 Les Ulis, France; Centre de Recherche en Cancérologie de Marseille, Aix Marseille Université, Inserm, CNRS, Institut Paoli-Calmettes, Equipe Leader Fondation ARC 2021, 13009, Marseille, France

**Keywords:** Botulinum toxin, neurons, SV2, BoNT/A, translocation, SNAP25

## Abstract

Botulinum neurotoxin A (BoNT/A) is a highly potent proteolytic toxin specific for neurons with numerous clinical and cosmetic uses. After uptake at the synapse, the protein is proposed to translocate from synaptic vesicles to cytosol through a self-formed channel. Surprisingly, we found that after intoxication proteolysis of a fluorescent reporter occurs in the neuron soma first and then centrifugally in neurites. To investigate the molecular mechanisms at play, we use a genome-wide siRNA screen in genetically engineered neurons and identify over three hundred genes. An organelle-specific split-mNG complementation indicates BoNT/A traffic from the synapse to the soma-localised Golgi in a retromer dependent fashion. The toxin then moves to the ER and appears to require the Sec61 complex for retro-translocation to the cytosol. Our study identifies genes and trafficking processes hijacked by BoNT/A, revealing a complex route for efficient intoxication that contradicts the currently accepted model of BonT intoxication.

## Introduction

Botulinum toxins (BoNTs) are the most potent toxins known, with a lethal dose of less than a microgram; paradoxically they are also one of the most widely used drugs today for neuromuscular conditions and various aesthetic procedures. BoNTs block overactive muscles involved in neuromuscular disorders or wrinkles by preventing nerve stimulation. The high potency of BoNTs allows for precise localized treatments through injection of small quantities of toxins. Their long-lasting effects make them attractive treatments despite the required injection. These remarkable properties of BoNTs are derived from their unique intoxication biology. Yet, key features of this biology remain poorly understood.

BoNTs target synapses at the neuro-muscular junction, where they cleave specific cytosolic proteins involved in the release of the acetylcholine neurotransmitter in neurons (Dong et al., 2019; Montecucco and Schiavo, 1994). As BoNTs are proteins of ∼150kDa, they cannot diffuse through cellular membranes. The key biological features of BoNTs are: 1) high binding specificity for neurons, specifically neuronal synapses, 2) ability to reach the cytosol of neurons, 3) high specificity for a molecular target once in the cytosol. Of these three features, the second is probably the least understood.

BoNTs are naturally expressed by the gram-positive bacterium *Clostridium botulinum* (*C. botulinum*) as a single ∼150kDa polypeptide chain, then post-translationally cleaved to produce a 50kDa N-terminal catalytic light (L) chain, which is a zinc metalloprotease and a 100kDa C-terminal heavy (H) chain. The L and H chains are linked via a disulfide bond and other noncovalent interactions to form the active toxin. *C. botulinum* produces seven serotypes of BoNTs, A to G, and while they all interfere with neurotransmission, they have different cell surface receptors and, interestingly, different kinetics of action across subtypes (Pellett et al., 2015b; Rasetti-Escargueil and Popoff, 2020).

BoNT/A is the most clinically-used serotype, it binds to the synaptic membrane of neurons with high specificity, with synergistic binding to the synaptic vesicle protein 2 (SV2) and the trisialogangloside GT1b (Dong et al., 2006; Yowler and Schengrund, 2004). After internalization and translocation to the cytosol, the light chain of BoNT/A binds and cleaves the SNARE (Soluble N-ethylmaleimide-sensitive factor Attachment protein REceptor) SNAP25 at the Q167R residue. As SNAP25 is essential for the release of synaptic vesicles, its inactivation results in the inability of neurons to release acetylcholine, the key neurotransmitter at neuromuscular junctions (Dong et al., 2019; Montecucco and Schiavo, 1994). As the toxin proteolytic activity is highly specific, the neuron remains viable and able to recover once the toxin has been cleared (Peng et al., 2013).

The current model for intoxication postulates cell surface binding and internalisation in endosomes, then, due to changes of pH there are conformational changes in the heavy chain that leads to the formation of a channel which ushers the L chain into the cytosol of the axonal bouton (Azarnia Tehran et al., 2017; Pirazzini et al., 2014). The heavy chain is proposed to contain an N-terminal translocation domain (HN) in addition to the C-terminal receptor-binding domain (HC). Once or while the L chain is translocated, it is released from the H chain by disulfide bond reduction, mediated by the thioredoxin reductase 1 (TXNRD1) (Dong et al., 2019; Pirazzini et al., 2013; Rossetto et al., 2021). After refolding of the L chain, it binds and cleaves SNAP25 at a specific residue.

Parts of this scenario have been challenged by studies showing that there are no distinct structural changes to the H and L domains upon acidification and by the difficulty in observing in vitro a self-translocation process (Araye et al., 2016; Galloux et al., 2008). Studies have also shown that BoNT/A is trafficked beyond the axon boutons in non-acidic and non-recycling vesicles (Antonucci et al., 2008; Harper et al., 2016; Restani et al., 2012). In addition, the current model fails to explain the delayed onset of action, with hours to days between the time of injection and muscle paralysis (Ledda et al., 2022). Arguably, the main problem of the model is its failure to propose a thermodynamically consistent explanation for the directional translocation of a polypeptidic chain across a biologial membrane. Other known instances of polypeptide membrane translocation such as the co-translational translocation into the ER indicate that it is an unfavorable process, which requires cellular energy (Alder and Theg, 2003).

Other similar toxins, like the Cholera toxin, Ricin and Pseudomonas exotoxin A, follow a complex intracellular trafficking route, first from endosomes to the Golgi apparatus, then to the endoplasmic reticulum where they translocate across the membrane. The translocation event itself relies on the host translocon machinery or other ER endogenous complexes (Moreau et al., 2011; Nowakowska-Gołacka et al., 2019; Zhang et al., 2013).

In this study, we present a novel assay based on genetically modified human progenitors able to generate functional neurons (Donato et al., 2007; Song et al., 2019). The reporter assesses BoNT/A proteolytic activity in live cells and is amenable to a high throughput screening. Complete cleavage of the reporter requires nearly 72h of exposure. Surprisingly, we find that BoNT/A proteolytic activity is first detected in the soma of neurons, 24 to 48 hours before reaching the end of neurites. To investigate the underlying reasons, we performed a genome-wide RNAi survey of the host factors required for BoNT/A intoxication. Our results reveal a high number of genes linked to membrane trafficking, suggesting that BoNT/A follows a complex intracellular route. We use another set of reporters, based on GFP reconstitution to retro-axonal traffic is required for BoNT/A to reach the ER, where it translocates to the cytosol using the Sec61 translocon. These findings help explain the delayed effect of the toxin and could pave the way to improved therapeutics.

## Results

### ReD SNAPR: neuronal cells expressing a BoNT/A reporter derived from SNAP25

To establish a high-throughput assay for BoNT/A activity, we selected an abundant and consistent source of neuronal cells, the ReNcell VM, a v-myc-transformed human neuronal stem cell line. This cell line has the ability to differentiate into neurons in about two weeks after withdrawal of EGF and bFGF from the culture medium. Over this period, neurites are formed and the neuronal markers, beta3-tubulin and MAP2 increase significantly in differentiated cells **(Supplementary Figure 1A)**. In parallel, the low levels of the oligodendrocyte marker CNPase expressed in the stem cells are further diminished.

To detect BoNT/A activity, we generated a chimeric reporter protein composed of SNAP25 flanked by the red fluorescent protein called tagRFPT (tRFPT) and the green fluorescent protein tagGFP (tGFP) at its N-terminus and C-terminus respectively (Shaner et al., 2008). The construct was named SNAPR (<u>SNAP</u>25 tagged with <u>R</u>FP and GFP). Using lentiviral transduction, we generated ReNcell VM cells stably expressing SNAPR (**Figure 1A**). We coined this cell line Red-SNAPR for <u>Re</u>Ncell-<u>d</u>erived, expressing <u>SNAPR</u>. After BoNT/A cleaves the SNAP25 moiety at Q197R, the C-terminal tGFP-containing moiety is rapidly degraded while the rest of the construct is preserved (**Figure 1A)**.

**Figure 1.**
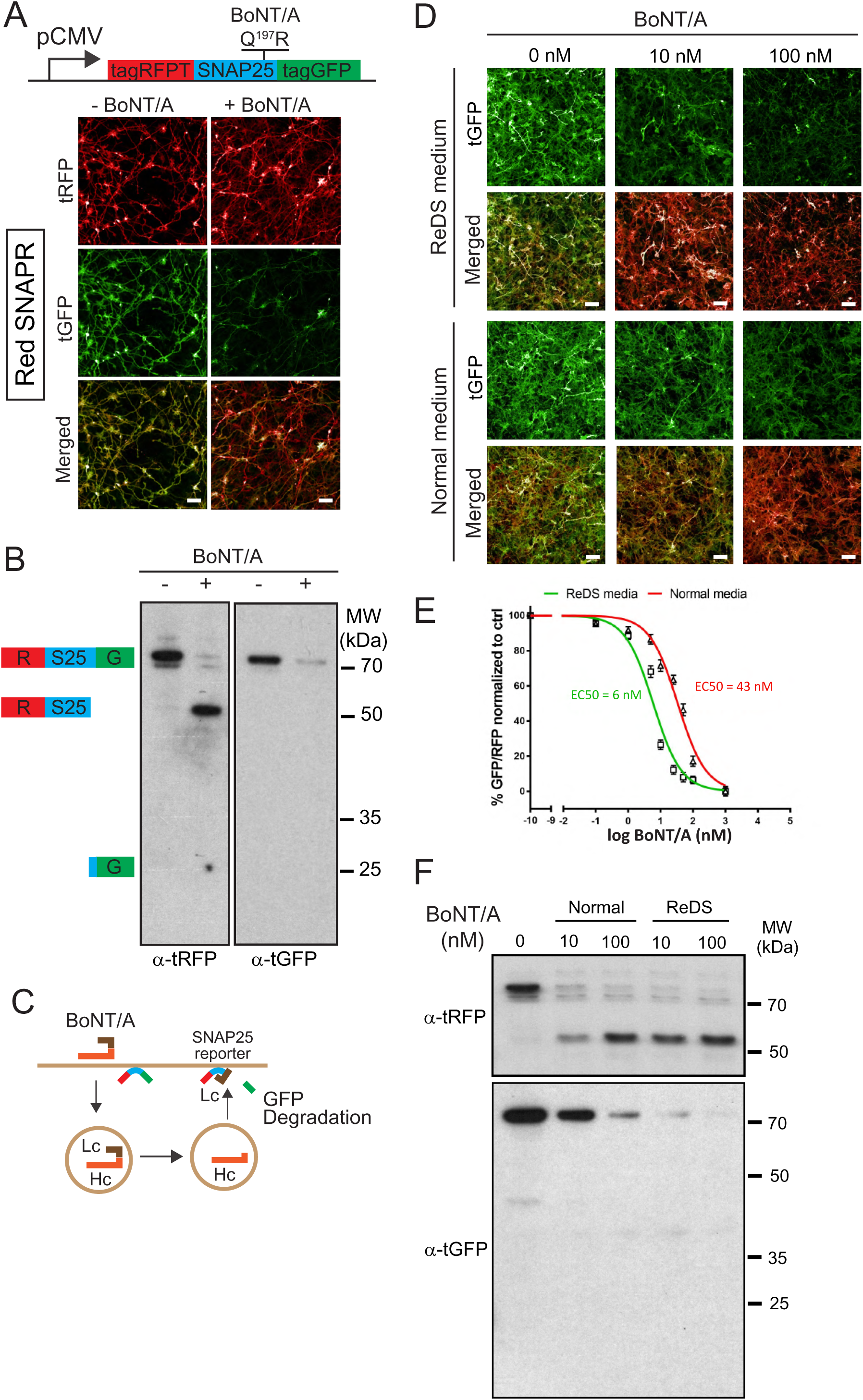
Differentiated BoNT/A reporter cell line, Red SNAPR, is highly sensitive to BoNT/A intoxication. **(A)** Schematic diagram of the BoNT/A reporter construct, SNAPR. Representative images of ReNcell VM cell line, Red SNAPR, stably expressing SNAPR and incubated with 100nM BoNT/A for 48 hours. **(B)** Western blot of cell lysates from **(A)** probed with tRFPT and tGFP antibodies. **(C)** Schematic diagram of the BoNT/A intoxication assay. **(D)** Red SNAPR differentiated in normal or ReDS medium, then incubated with 0, 10 and 100 nM BoNT/A for 48 hours. Quantification of EC_50_ dose response of BoNT/A in ReD SNAPR cells incubated with 0 - 100nM BoNT/A in normal and ReDS medium using GFP/RFP ratio readout. **(E)** Western blot of cell lysates from **(D)** probed with tRFPT and tGFP antibodies. Scale bars 50 μm.

After incubation with 100 nM BoNT/A for 48 hours, tGFP fluorescence was noticeably diminished compared to tRFP, whose signal remains stable. By western blot, a ∼75 kDa SNAPR construct was detected by both tRFP and tGFP antibodies, and cleaved to a 50 kDa product only detected by tRFP antibody, while the expected ∼25 kDa tGFP fragment was undetectable (**Figure 1B)**. This stable cell line has unaltered differentiation potential (**Supplementary Figure 1B).**

To test whether BoNT/A intoxication was dependent on neuronal activity, we used the neurotrophic factors GDNF and BDNF to enhance neuronal differentiation. We further supplemented the medium with high salt (KCl and CaCl) for neuronal stimulation (Harper et al., 2011; Pellett et al., 2015a). This differentiation and stimulation media (ReDS media) resulted in an improvement of sensitivity by an order of magnitude in the imaging assay (**Figure 1D, E).** This increased sensitivity was confirmed by western blot in which the EC_50_ of BoNT/A intoxication improved from 43 nM to 6 nM (**Figure 1F**). This strongly suggests that toxin binding and uptake occurs at the level of active synapses, as is the case in vivo.

### BonT/A activity is first detected in the soma of neurons

During the initial experiments, we noticed that optimal degradation of the reporter required 48h after adding the toxin. This delay was surprising considering the model of rapid translocation after internalisation. As the reporter allows live detection of BonT/A proteolytic activity, we were curious to observe the distribution of BonT/A activity within the neurons over time. We thus imaged cells at 24, 48 and 72hr after intoxication. To facilitate imaging of individual neurites, a co-culture of Red-SNAPR to ReNcell VM cells (1:4 ratio) was implemented (**Figure 2A).** Surprisingly, at 24 hours, there was no loss of GFP signal in the terminal part of neurites but visible degradation at the neurite hillock. At 36 hours, tGFP degradation progressed towards the neurite terminals and most of the tGFP signal was eliminated by 48 hours. The pattern suggests a slow distribution of BoNT/A from the cell body to the terminus of neurites (i.e. ∼16µm/h) **(Figure 2B)**.

**Figure 2:**
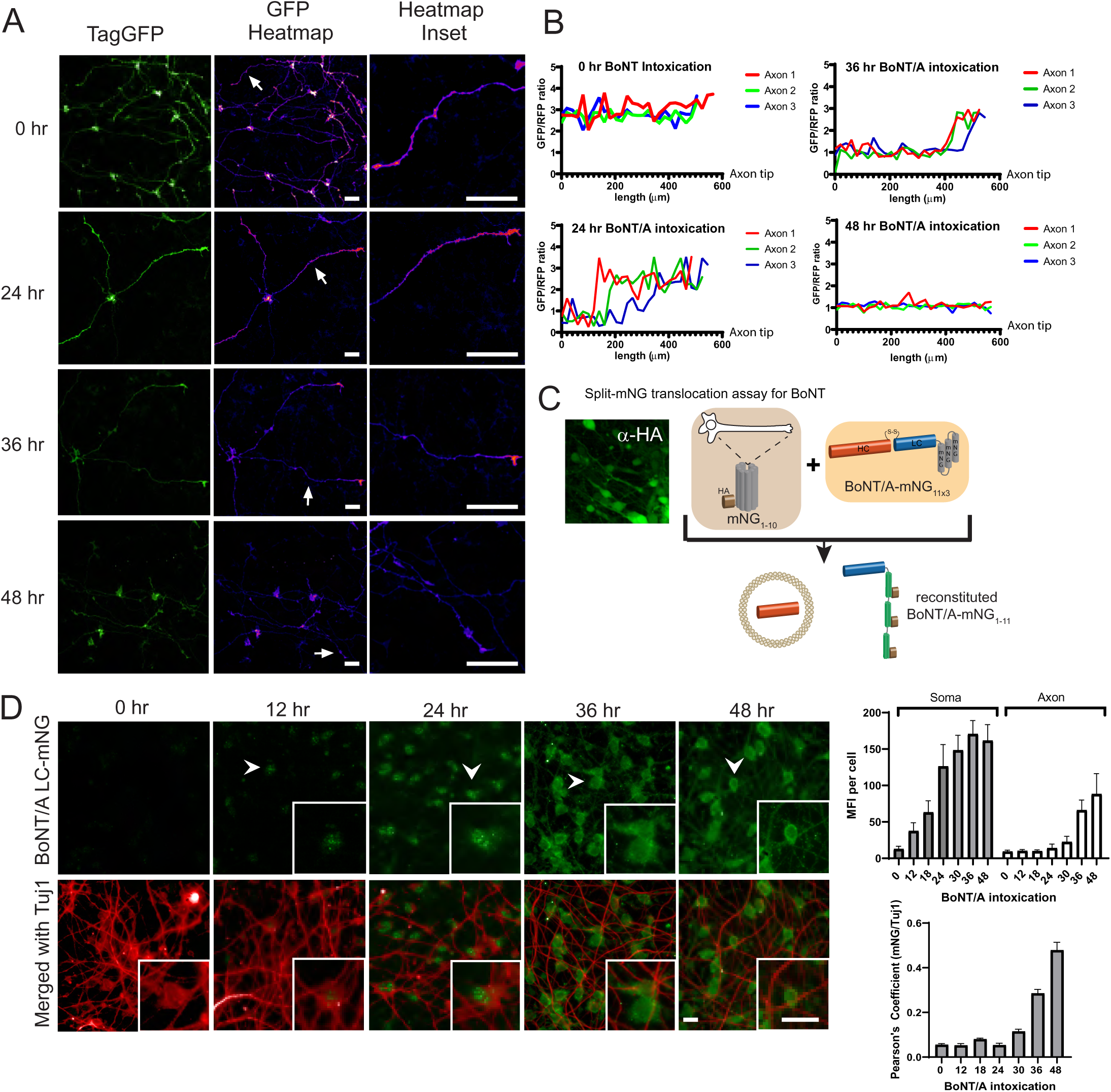
BoNT is first detected in the neuronal soma then emanate to the axons. **(A)** Red SNAPR cells co-cultivated at ¼ with ¾ of unlabeled Ren-VM were imaged for 48h after addition of BoNT/A. The GFP color-coded intensity signal is displayed in the second column. **(B)** Quantification of GFP/RFP signal along the length of individual neurites (Axon 1,2,3) at various time points. **(C)** Schematic diagram of split-mNG (NeonGreen Fluorescent Protein) detection system, consisting of ReNcell VM expressing cytosolic mNG_1-10_ and mNG_11_-tagged BoNT/A. Fluorescence occurs after binding of mNG_11_-tagged BoNT/A to mNG_1-10_. **(D)** Time-course of fluorescence reconstitution after exposure to mNG_11_-tagged BoNT/A and quantification of Mean Fluorescence Intensity in the soma and neurites of cells. Scale bars 20 μm.

### BoNT/A protein is also detected first in neuronal soma

SNAPR reports on BoNT activity and it is conceivable that BoNT/A might not be active immediately after translocation, thus potentially affecting the spatiotemporal pattern of proteolytic activity. To directly detect BoNT protein in the cytosol, we generated a ReNcell VM cell line expressing an HA-tagged split monomeric NeonGreen (mNG) protein targeted to the cytosol (Cyt-mNG_1-10_) (Feng et al., 2017). We verified that Cyt-mNG_1-10_ was expressed using the HA tag, the expression was homogeneously distributed in differentiated neurons and we observed no GFP signal (**Figure 2C**). We also generated and produced the complementary mNG_11_ fused to BoNT/A (BoNT/A-mNG_11_) with three beta-strands of mNG in tandem (**Figure 2C)**.

We next imaged the neurons after different intoxication times (**Figure 2D**). After 12h, the first signal was observed in the cell body of neurons, in concordance with the pattern of BoNT activity. The fluorescence appeared in specks first, then converted to a more diffuse pattern. At 24h and 36h, the GFP pattern became more diffuse and started to spread in the neurites. The initial pattern might reflect reconstitution of GFP proteins at the site of extrusion from membranes. By 48h, the GFP signal had filled up the neurites, indicating that BoNT/A had diffused throughout the cell (**Figure 2D**). Quantification of the GFP signal confirmed an initial accumulation in the soma followed by an increase in neurites (**Figure 2D**). The late (>24h) and gradual accumulation in neurites was further confirmed by quantification of intensity correlation between BoNT/A LC-mNG and α-Tuj1, a neuronal marker enriched in neurites (**Figure 2D**).

Altogether, the data indicates that BoNT/A translocates into the cytosol at the level of the soma. Based on the dependency of the toxin on neuronal activity and current knowledge of BoNT/A receptors, it is likely that BoNT/A is internalised at the tip of neurites, where synapses form. Thus, the data would suggest that BoNT/A requires retrograde trafficking before it can translocate.

### A genome-wide RNAi screen reveals numerous positive and negative regulators of BoNT intoxication

To elucidate the molecular mechanisms required for BoNT/A intoxication and understand the surprising pattern of appearance, we decided to systematically survey the genes required for BoNT/A translocation.

We first optimized the liposome-mediated delivery of siRNA in the Red-SNAPR cells (**Supplementary Figure 1C).** As a positive control, we targeted TXNRD1 as a factor known to be required. Cells differentiated for 2 weeks were transfected with TXNRD1 siRNA for 3 to 5 days and analysed by image analysis. Undifferentiated and differentiated ReNcell VM displayed ∼70% knockdown compared to non-targeting control as measured by image analysis of TXNRD1 staining. We also targeted SNAPR using an siRNA targeting SNAP25. This approach achieved an 80% reduction in signal (**Supplementary Figure 1D**).

As neurons cannot be passaged after differentiation, a forward siRNA transfection pipeline was developed (**Figure 3A**). The genome-wide screen was carried out in duplicate using pools of 4 siRNAs per gene and targeting 21,121 human genes in total. Differentiated Red-SNAPR cells on laminin-coated 384-well imaging plates were incubated with siRNA complexes for 3 days and intoxicated with BoNT/A for 2 days before imaging. The positive control siTXNRD1 rescued the tGFP signal reproducibly (**Figure 3B**).

**Figure 3.**
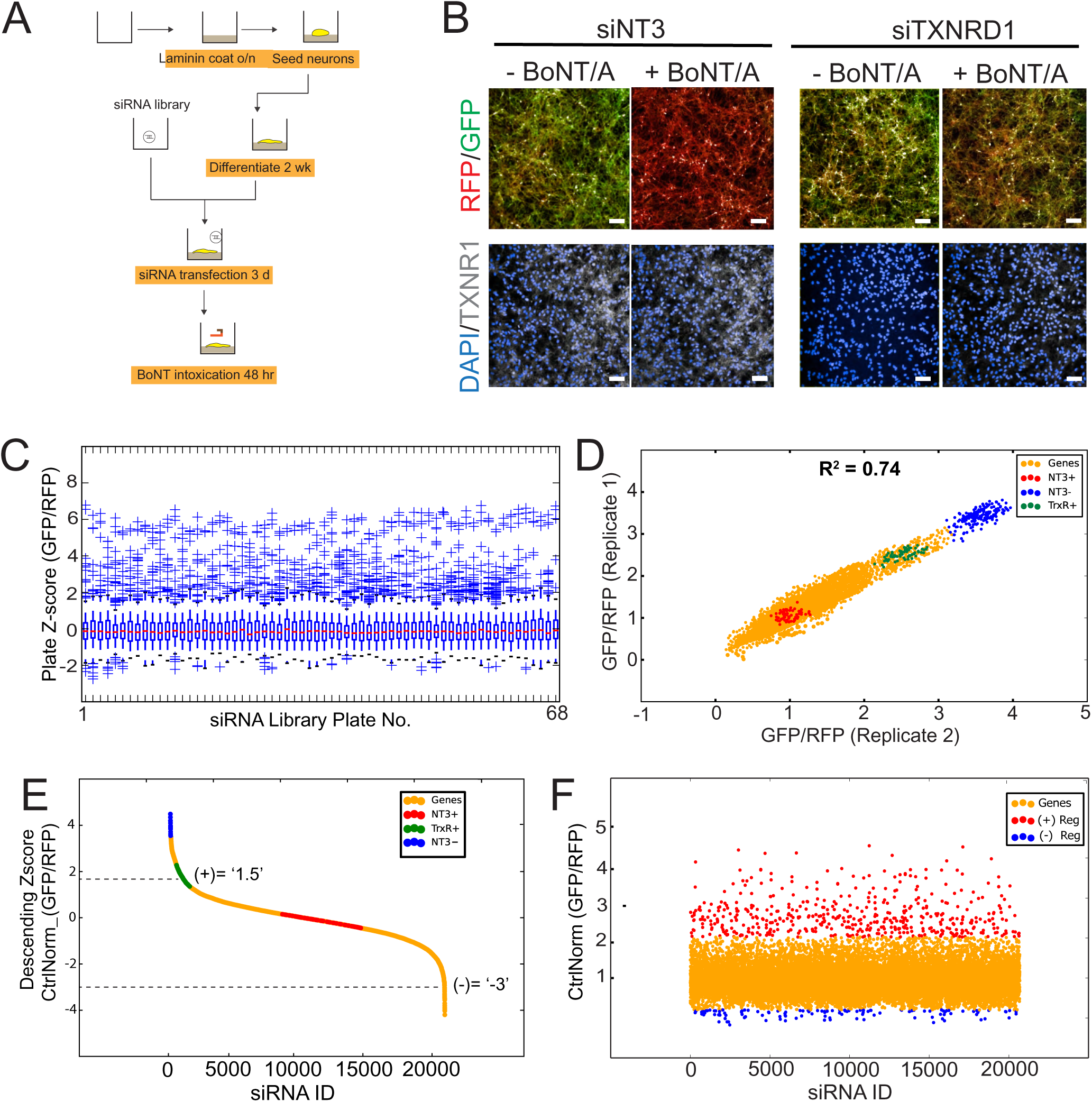
Genome-wide RNAi screen of BoNT/A-treated Red SNAPR cell line. **(A)** Schematic diagram of assay pipeline. **(B)** Cells treated with non-targeting control siRNA (siNT3) and positive control siRNA (siTXNRD1), then incubated with BoNT/A for 48 hours. Cells were stained with DAPI and TXNRD1 antibody post-fixation. Scale bars 50 μm. **(C)** ScreenSifter Z-score analysis of GFP/RFP ratio index of whole genome siRNA library plates. **(D)** R-squared analysis of both genome-wide RNAi screen replicates. Blue dots = No toxin control, Red dots = Toxin control, Green dots = Positive control (siTXNRD1). **(E)** Descending Z-score of control-normalized and averaged GFP/RFP ratio of genome-wide screen to determine cut-off values of +1.5 for positive regulators and −3 for negative regulators. **(F)** Chronological order of genome-wide control-normalized screen reflecting the positive regulators (red) and negative regulators (blue) derived from **(E)**.

We analyzed the two replicates of the whole genome-wide screen using the ScreenSifter software (Kumar et al., 2013) **(Supplementary Figure 2A)**. The data was converted to average plate Z-score, revealing low plate-to-plate variations (**Figure 3C)**. The controls such as the BoNT/A-treated, siNT3 (NT3+), non BoNT/A-treated siNT3 (NT3-) and BoNT/A-treated, siTXNRD1 (TXNRD+ in figure) were tightly grouped (**Figure 3D**). The assay had a robust Z-factor of 0.81. The two replicates were correlated with an R-value of 0.74. Genome-wide plots for individual replicates are shown in **Supplementary Figure 2B**. The tight clustering of the controls demonstrates high reproducibility between experiments. The data was ranked to establish cut-offs for hits selection (**Figure 3E)**.

Most hits (363) were genes required for BoNT/A intoxication (red dots); interestingly a significant fraction of hits (76) resulted in enhanced intoxication (blue dots) (**Figure 3F**). Many of the positive regulators resulted in higher rescue than the siTXNRD1 control, suggesting that previously unidentified molecular processes are critical for BoNT/A intoxication.

To exclude potential indirect effects, we first used nuclei counts and identified 289 genes that significantly affect neuronal survival (**Supplementary Figure 2C)**. 80 positive and 8 negative genes were sifted out of the hit list (**Supplementary Figure 2D**). Next, we carried out a duplicate deconvoluted siRNA screen on the hit list to exclude potential off-target siRNAs (Jackson and Linsley, 2010) (**Supplementary Figure 2E**). Using this approach, 35 genes could not be confirmed by two independent siRNAs and thus were sifted out. To ensure the hit list only contains genes significantly expressed in neurons, we carried out RNA sequencing analysis on differentiated Red SNAPR cells (**Supplementary Figure 2F)**. A cut-off threshold of counts-per-million (CPM) at 0.25 (log_2_CPM = −2) was used (grey bars) and 31 more genes (18 positive, 13 negative) were excluded.

### The surface expression of BoNT/A receptor, SV2, is highly regulated

We next focused on genes influencing BoNT/A cell surface binding by studying the surface expression of SV2, the BoNT/A receptor. We incubated fixed and non-permeabilized cells with an antibody against the extracellular domain of SV2A to quantify cell surface exposure (**Figure 4A**). Depletion of VAMP2, a known regulator of synaptic vesicle fusion and SV2 trafficking (Pennuto et al., 2003), resulted in 60% reduction of surface SV2 levels relative to non-targeting control (siNT3) (**Figure 4B**). By contrast, siTXNRD1 did not affect SV2 surface signal (**Figure 4B**).

**Figure 4.**
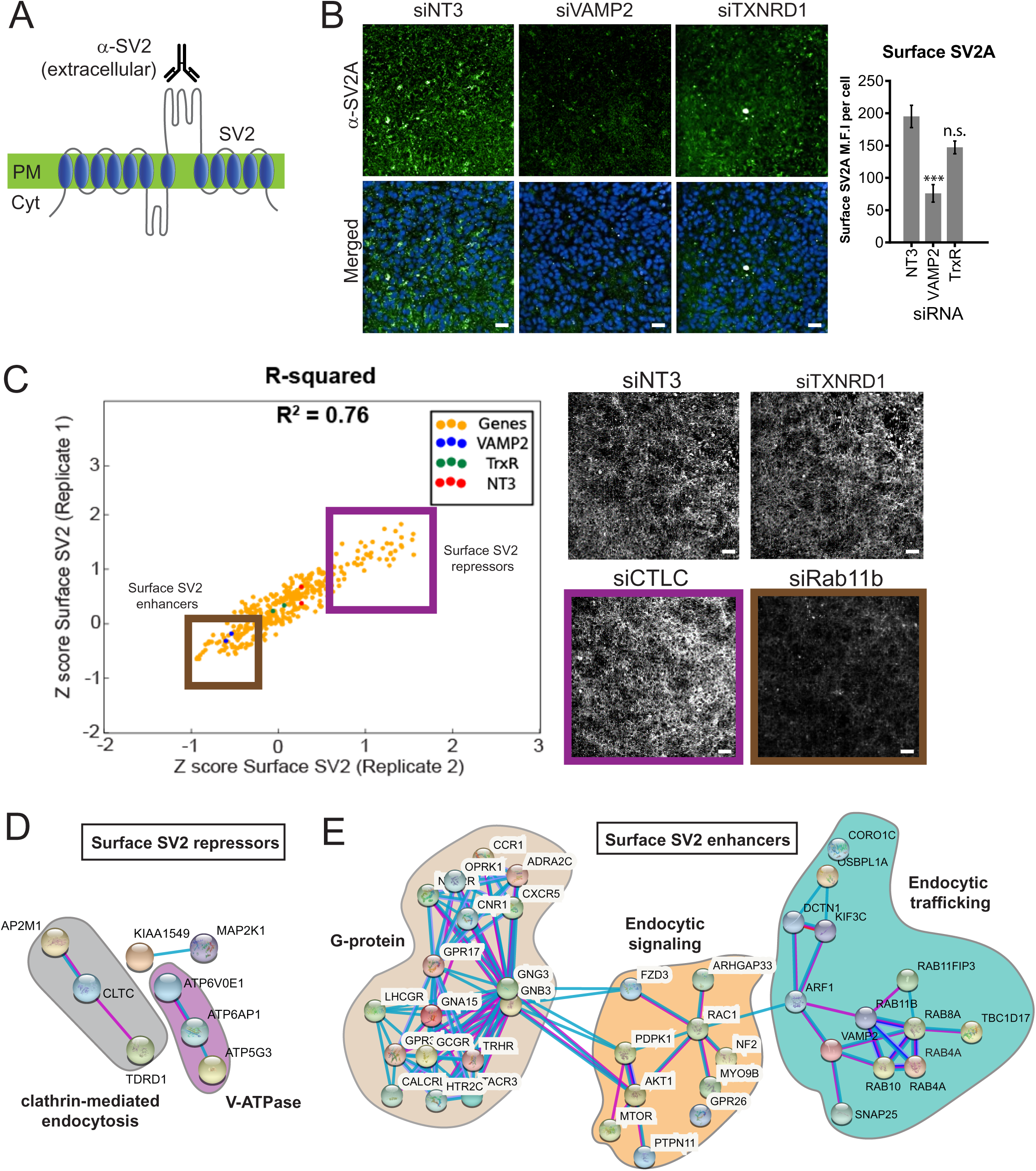
Surface expression of the BoNT/A receptor SV2 is regulated by endocytic trafficking and signaling genes, forming a cohort of the positive regulators of the genome-wide screen. **(A)** Detecting surface SV2 using a specific antibody against the extracellular loop of SV2. **(B)** siVAMP2 as a positive control for surface SV2 expression. ReNcell VM treated with siNT3, siVAMP2 and siTXNRD1 for 3 days and stained with SV2A antibody after fixation without membrane permeabilization. Quantification of surface SV2 per cell where whole image SV2A MFI was divided by total nuclei count (DAPI) from 3 fields of view and 3 independent replicates. Scale bars 50 μm. **(C)** Replicate screen results for surface SV2 regulators from genome-wide positive hits. Purple box = surface SV2 repressors e.g. siCTLC (clathrin light chain); Brown box = surface SV2 enhancers e.g. siRab11b. siNT3 and siTXNRD1 does not regulate surface levels of SV2. Scale bars 50 μm. **(D)** STRINGS analysis of surface SV2 repressors. **(E)** STRINGS analysis of surface SV2 enhancers.

We screened a library of hits identified in **Figure 3F** and identified 105 genes that affect SV2, with 38 gene depletions reducing (Repressors, SV2(-) in table) and 67 gene depletions enhancing (Enhancers, SV2(+) in table) surface SV2 (**Figure 4C, Supplementary Table 1)**.

For instance, depletion of clathrin light chain (siCTLC), a known regulator of endocytosis, increased surface staining of SV2 (Yao et al., 2010). By contrast, depletion of Rab11b decreased surface SV2 which could be due to a block in recycling of SV2 from endosomes to the cell surface (Giorgini and Steinert, 2013). Using STRINGS analysis on ‘surface SV2 repressors’, we identified a closely-associated network of genes related to clathrin-mediated endocytosis such as AP2M1, CTLC and TDRD1. In addition, distinct subunits of the V-ATPase were identified which can function as adaptins to facilitate endocytosis (Geyer et al., 2002) (**Figure 4D**). The majority of the surface SV2 enhancers are associated with G-protein signaling, which controls neuronal excitation (**Figure 4E)**. This agrees with earlier findings (Figure 1D) where increased neuronal activity favors intoxication as observed when K^+^/Ca^+^ are spiked in the medium. Endocytic signaling and membrane trafficking gene families such as Rabs, Arfs and SNAREs were also revealed, congruent with a role in SV2 exocytosis (**Figure 4E**).

### Network analysis reveals regulators of signaling, membrane trafficking and thioreductase redox state involved in BoNT/A intoxication

Among the positive regulators of the screen, 135 hits did not influence significantly surface SV2 levels and are thus likely to function in post-endocytic processes (**Supplementary Table 2**). However, we cannot formerly exclude that they could affect binding of BonT to the cell surface independently of SV2. 92 positive regulators (required for intoxication, in red) and 43 negative regulators (reducing intoxication, blue) were mapped to their intracellular localities (**Figure 5A)**. Several gene products were localized to endosomes and 16 were associated with the Golgi and ER. At the Golgi, one hit was the glycosylation enzyme B4GALT4, known to be involved in the biosynthesis of the ganglioside co-receptor of BoNT/A, G_T1b_. Using the STRING database, a protein-protein interaction network of hits was generated (**Figure 5B)** (Szklarczyk et al., 2021). A subnetwork was constituted of heat-shock protein (HSP) chaperones of the HSP70 family (HSPA4 and HSPA14). Another chaperone, HSP90, has been linked to the translocation of the clostridial toxins C2 toxin and BoNT/A into the cytosol (Azarnia Tehran et al., 2017; Haug et al., 2003). As HSP90 and HSP70 act in a sequential cascade for protein folding (Genest et al., 2019), our results suggest that such a chaperone cascade might help BoNT/A LC refold in the cytosol after translocation.

**Figure 5.**
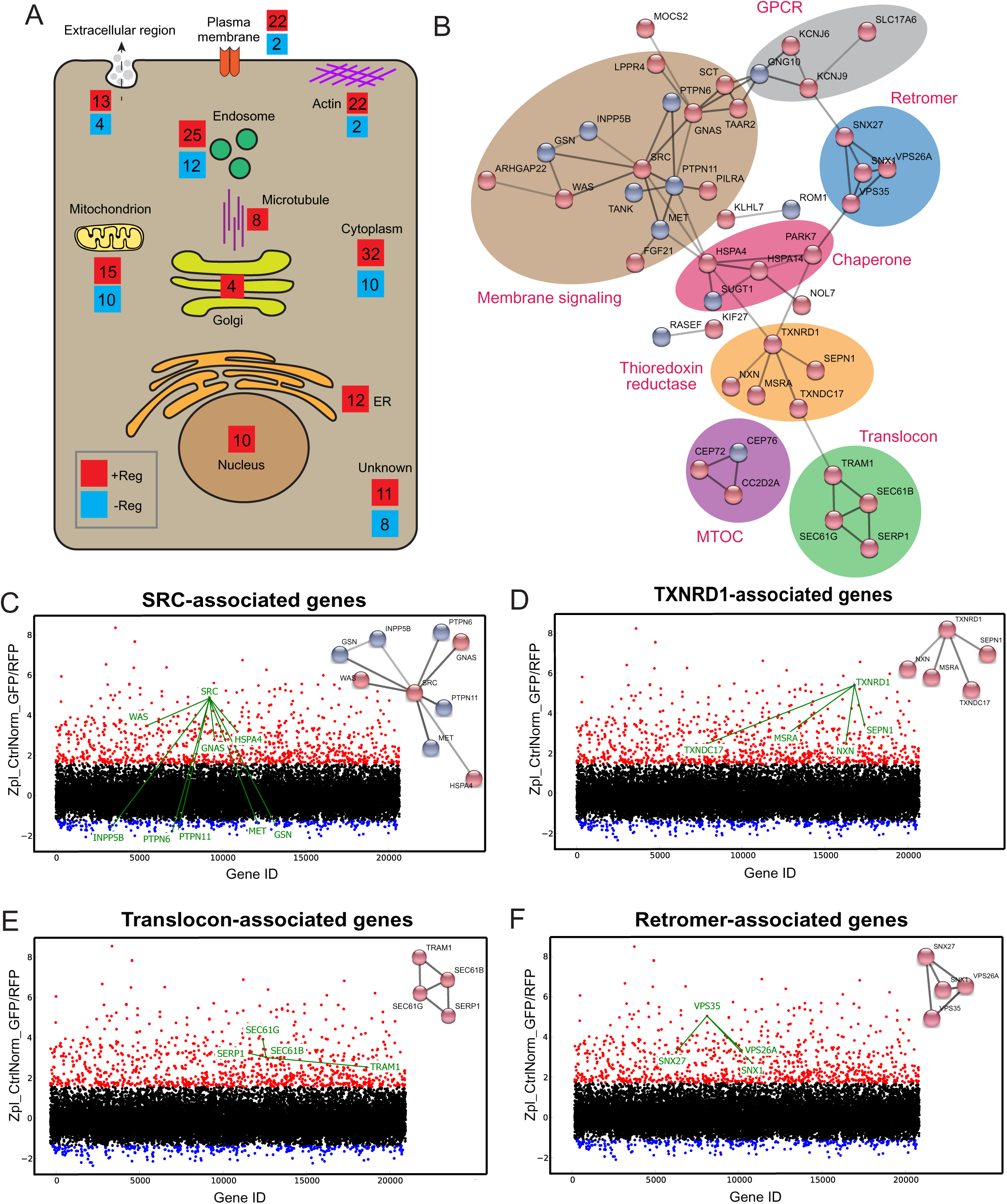
Protein network analysis of genome-wide hits reveals known and novel regulators of BoNT/A intoxication. **(A)** Summary diagram of all positive and negative hits mapped to their intracellular localities. **(B)** STRINGS network analysis of connected hits (non-connected hits are excluded). Associated genes tied to respective cellular molecular complex/processes are bounded and annotated. ScreenSifter analysis revealed **(C)** Src-associated genes **(D)** TXNRD1-associated genes **(E)** Translocon-associated genes **(F)** Retromer-associated genes.

A large subnetwork of signaling and GPCR-related genes was identified, centered on the SRC tyrosine kinase (**Figure 5B**). Depletion of SRC resulted in one of the most stringent intoxication blocks (**Figure 5C**). Interestingly, the tyrosine phosphatase PTPN6 was also one of the strongest negative regulators (Chong and Maiese, 2007) (**Figure 5C**). It has been proposed that SRC directly phosphorylates and regulates BoNT LC catalytic domain (Ferrer-Montiel et al., 1996; Ibañez et al., 2004; Kiris et al., 2015). On the other hand, SRC and its partners are crucial for retrograde membrane trafficking and might influence post-endocytic BoNT/A trafficking (Chia et al., 2021; Sandilands and Frame, 2008). The screen identified TXNRD1, consistent with previous literature, but also several genes involved in thioredoxin reduction, including Methionine sulfoxide reductase A (MsrA), Thioredoxin domain-containing protein 17 (TXNDC17), Nucleoredoxin (NXN) and Selenoprotein N (SEPN1) (**Figure 5D**). It suggests that these proteins either interact directly with BoNT/A or are required for TXNRD1 function.

A set of genes linked to the translocon (SEC61G, SEC61B, TRAM1, SERP1) were identified (**Figure 5E**). GO:MF analysis (Molecular Function) indicates high enrichment level of these ER translocon genes **(Supplementary Figure 3A).** The requirement of the translocon suggests its involvement in translocation of the toxin to the cytosol as is the case for other toxins (Moreau et al., 2011; Nowakowska-Gołacka et al., 2019). However, this hypothesis implies that the toxin travels from endocytic vesicles at its site of internalisation to ER membranes where the translocon is localized.

Interestingly, genes for the retromer were also identified, VPS35, VPS26A, SNX1 and SNX27, with a high level of enrichment in gene ontology analysis **(Figure 5F, Supplementary Figure 3B).** These genes did not significantly affect surface SV2 levels. The retromer has been linked to endosomes to Golgi traffic, suggesting the toxin might need to traffic between these organelles.

### Retro-axonal traffic of the BoNT/A receptor SV2 requires the retromer

We next analysed the effect of VPS35 depletion on the kinetics of BoNT/A-mNG_11_ arrival in the cytosol. The control siNT3 treated cells showed a progressive increase in signal from 12 to 48h, with a progressive appearance of signal in the neurites (**Supplementary Figure 4A)**. By contrast, siVPS35 treated cells only displayed some soma-localised signal after 48h (**Supplementary Figure 4A).**

To test directly the trafficking of BoNT/A intracellularly proved difficult as reagents we tested were not sensitive enough to image the trafficking of the toxin. Since BoNT/A binds to SV2 at the cell surface, we wondered whether the receptor itself was trafficked retrogradely in neurites. We generated a ReNcell VM cell line expressing a chimera of SV2 with the Dendra fluorescent protein. We found that Dendra-SV2 is distributed in the whole neuron, similarly to the endogenous protein (**Supplementary Figure 4B)**.

Next, we depleted cells of VPS35 and found that Dendra-SV2 accumulated in bulbous structures at the neurite tips. To quantify this phenomenon, we compared the number and size of SV2 puncta in neurites in control and siVPS35-treated cells. Using ImageJ to threshold and select for particles of interest, we measured the number of particles along every 50μm segment of the neurite, starting from the neurite tips **(Supplementary Figure 4C)**. In control cells, SV2 punctas were homogeneously spread throughout the neurites. However in VPS35-depleted cells, SV2 punctas were enriched at the neurite tip (1-50μm) and were significantly reduced with the rest of the neurite. These puncta were also much larger than in control cells. This evidence indicates that SV2 is trafficked retro-axonally in a retromer-dependent fashion, thus consistent with the notion of BoNT/A retrogradally to the neuronal body bound to its receptor.

### BoNT/A trafficks through the Golgi apparatus

The implication of the retromer suggested that BoNT/A transits through the Golgi apparatus. To test this hypothesis, we sought a method to detect BoNT/A in different intracellular compartments. We decided to exploit the split-monomeric Neon Green (mNG) fluorescence reconstitution approach (Luong et al., 2020). This approach relies on having the 10 beta strands of Neon Green fluorescent protein (mNG_1-10_) and the 11th beta strand in different proteins. When the two proteins can interact, fluorescence is reconstituted. We generated a ReNcell VM cell line stably expressing mNG_1-10_ fused with an HA tag and a fragment of β1,4-galactosyltransferase 1 for targeting to the Golgi (Golgi-mNG_1-10_) (Luong et al., 2020) (**Figure 6A**). We verified by immunofluorescence that the construct was strictly localized at the Golgi by the HA antibody staining and that no GFP fluorescence was detectable (**Figure 6B**).

**Figure 6.**
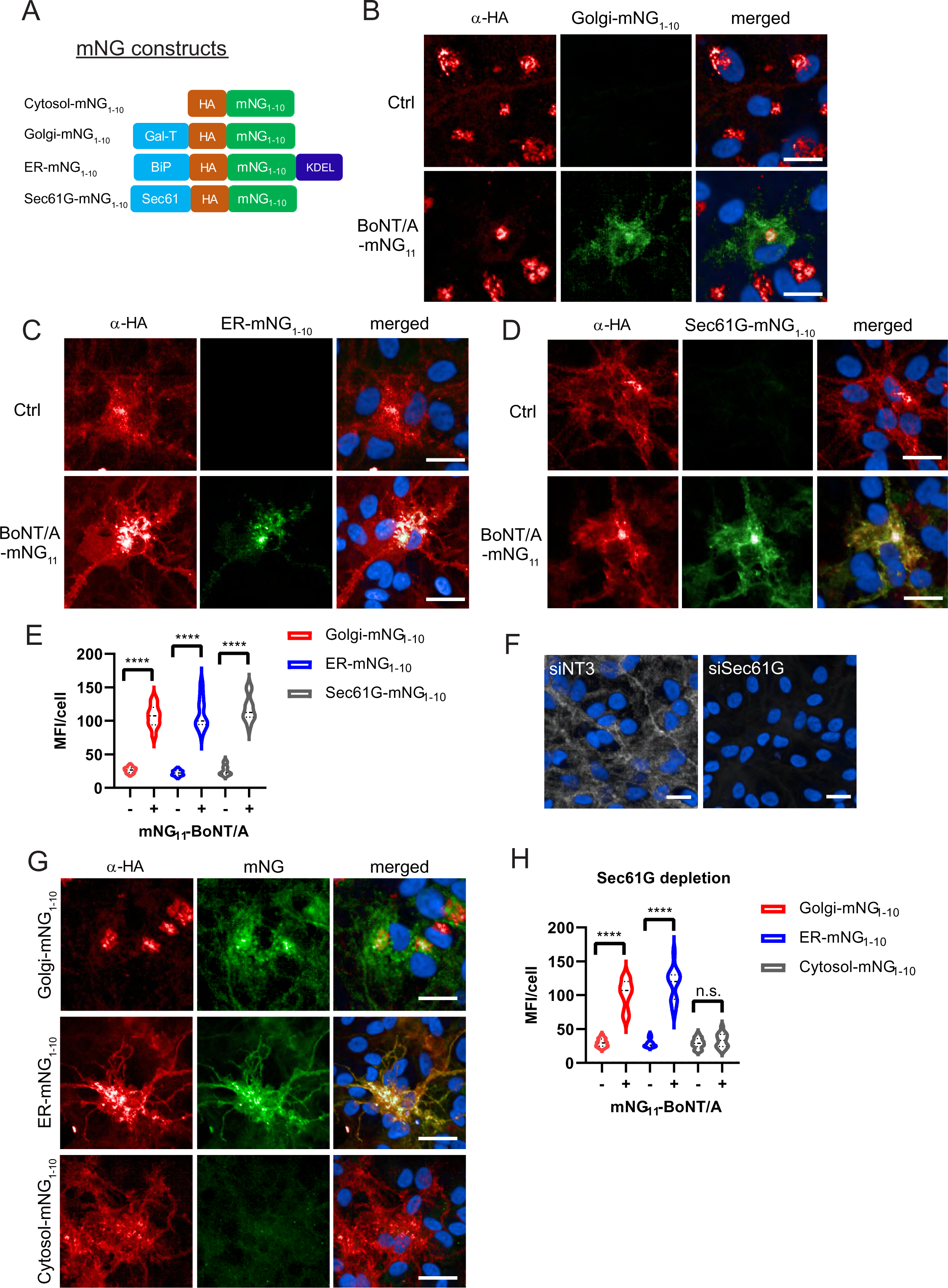
BoNT/A is retrogradely trafficked through Golgi and ER membranes as revealed by split-GFP reconstitution. (A) Split-mNG constructs targeted to cytosol, Golgi, ER lumen (KDEL sequence) and ER membranes (Sec61 transmembrane domain) were stably expressed in ReNcell VM and incubated with 50 nM of BoNT/A-mNG_11_. Control cells (without BoNT/A-mNG_11_) are shown at top panels of each condition **(B)** Cells expressing Golgi-mNG_1-10_ and showing reconstituted mNG fluorescence. Overlap of HA antibody and mNG signals in the Golgi **(C)** ER-mNG_1-10_ expressing cells showing mNG fluorescence and distinct overlap of HA antibody and mNG in the ER. **(D)** Sec61G-mNG_1-10_ expressing cells showing mNG fluorescence distinct overlap of HA antibody and Sec61G signals. **(E)** Quantification of MFI/cell from **B-D** showing increased mNG fluorescence in Golgi, ER and Sec61G-expressing cells after addition of BoNT/A-mNG11. **(F)** Representative image of SEC61G depletion in ReNcell VM. **(G)** Golgi-mNG_1-10_, ER-mNG_1-10_ and Cytosol-mNG_1-10_-expressing cells are depleted with siSEC61G and incubated with BoNT/A-mNG_11_ for 48 hours. **(H)** Quantification of MFI/cell from **G**. Graphs are obtained from at least 300 cells across 3 independent experiments. Scale bars 20 μm.

Upon incubation for 48 h with BoNT/A-mNG_11_, we observed reconstitution of mNG fluorescence at the Golgi **(Figure 6B)**. In addition, there was mNG fluorescence in the cell soma with a reticulated pattern, which could correspond to the ER. Interestingly, the HA pattern was also partially looking like ER in these cells. Thus the reconstituted mNG was not only restricted to the Golgi, which could be due to retrograde trafficking after complementation. Nonetheless, the Golgi pattern clearly indicates that BoNT/A traverses this organelle before translocation. In ReNcell VM cells, the Golgi apparatus is located almost exclusively in the soma of neurons.

### BoNT/A trafficks to the ER and translocate through Sec61 complex

To test whether BoNT/A can reach the ER, we next developed a chimera of mNG_1-10_ localized to the ER, based on the soluble, ER-resident BiP protein (ER-mNG_1-10_) (Luong et al., 2020). As for the Golgi construct, there was no fluorescence in the absence of BonT/A and the construct had the typical pattern of an ER protein (**Figure 6C**). 48h after intoxication with BoNT/A-mNG_11_, we could detect mNG in the ER (**Figure 6C**). To further confirm that BoNT/A can reach a Sec61-enriched compartment, we also developed ReNcell VM cell lines stably expressing a split-mNG_1-10_ construct fused with the transmembrane protein Sec61G (**Figure 6A**). As for the BiP based construct, the Sec61 staining pattern before addition of the toxin was consistent with the ER. After intoxication, the reconstituted mNG fluorescence had a typical ER pattern (**Figure 6D**). Quantification of fluorescence per cell confirmed that the three reporters were equally accessible to BoNT/A, indicating that the toxin traffics through the Golgi and ER in neurons **(Figure 6E)**.

Next, to test whether Sec61 is involved in BoNT/A LC translocation, Golgi-mNG_1-10_, ER-mNG_1-10_ and Cytosol-mNG_1-10_-expressing cells were depleted of Sec61G with siRNA **(Figure 6F)**. When incubated with BoNT/A-mNG_11_, the reconstituted mNG fluorescence appeared in Golgi-mNG_1-10_ and ER-mNG_1-10_ cells, while there was a striking reduction of fluorescence in cytosolic mNG_1-10_ expressing cells **(Figure 6G and 6H)**. These results indicate that loss of Sec61G does not prevent BoNT/A uptake nor its trafficking to Golgi and ER compartments, but is instrumental to the translocation of BoNT/A LC into the neuronal cytosol at the soma.

Overall, these reporters reveal the complexity of BoNT/A trafficking in neurons: after internalisation at active synapses in neurites, the toxin is transported retrogradely to the Golgi, then the ER before being translocated to the cytosol.

## DISCUSSION

To improve the understanding of BoNT/A intoxication, we designed the engineered BoNT reporter cell line, Red SNAPR. The line is easily maintained, highly sensitive to the toxin and the assay is a direct readout of GFP fluorescence. The assay encompasses all steps of BoNT/A intoxication from binding and endocytosis to light chain translocation and substrate cleavage. While FRET-based reporter assays of BoNT have been established, they tend to be limited by photobleaching and spectral bleedthrough from donor/acceptor fluorescence. Moreover, FRET efficiency is drastically reduced in fixed cells (Anikovsky et al., 2008; Dong et al., 2004). By contrast, SNAPR cleavage is quantified through the ratio of RFP to GFP signal, thanks to the instability of the GFP moiety after cleavage, probably due to the N-terminal arginine residue generated by BoNT/A cleavage at Q^197^R (Tasaki et al., 2012). As noted by one reviewer, the assay may be sensitive to perturbation in the general rate of protein degradation, a consideration to keep in mind when evaluating the results of large scale screens. On the other hand, this convenient and simple assay could replace animal-based assays that are used to measure BoNT potency. In addition, SNAPR could be implemented in transgenic mice to study the process of BoNT/A inhibition in vivo.

The sensitivity of the ReNcell VM cells to BoNT/A is highly enhanced by treatments that favor the formation of active synapses, consistent with the fact that BoNT/A binds the synaptic protein SV2, which only becomes exposed at the surface when neurons fire synaptic vesicles. Consistently, a hundred genes regulating BoNT/A intoxication (a quarter of the hits) actually impact cell surface exposure of SV2. The genes identified are involved in endocytic coupling, G protein-coupled receptors and other signaling genes. Thus, it seems highly likely that in our experimental model, BoNT/A is internalised at the tip of neurites (with axonal properties).

After binding to SV2 and internalisation, BoNT/A LC does not appear to translocate in the synaptic bouton as the reporter is unperturbed. The spatiotemporal pattern of BoNT/A activity and the cytosolic detection of the protein both indicate that the toxin appears in the soma of neurons about 12h after the start of incubation. Presumably in our cell culture system, these 12 hours are required for BoNT/A trafficking from synapses to the soma’s cytosol. Progression of cleavage of the SNAPR reporter from the soma to the neurite’s terminal requires an additional 36 h. The split-GFP organellar reporters indicate that BoNT/A traffics from endosomes to Golgi and then to the ER.

The genetic signature of BoNT/A intoxication requirements is consistent with this trafficking route. For instance, a significant positive regulator is the tyrosine kinase SRC. We and others have reported on the role of SRC at the Golgi/ER interface (Pulvirenti et al., 2008; Weller et al., 2010). We previously reported SRC’s role in regulating Pseudomonas Exotoxin A (PE) trafficking between Golgi and ER (Bard et al., 2003). More recently, we have shown the role of Src in directly controlling GBF1, a GTP Exchange Factorinvolved in Golgi to ER traffic (Chia et al., 2021). Interestingly, the counteracting tyrosine phosphatases PTPN6 and PTPN11 are negative regulators of BoNT/A and their depletion favors intoxication (Frank et al., 2004; Somani et al., 1997). To note, SRC has also been proposed to act directly on the toxin to activate it (Ferrer-Montiel et al., 1996; Ibañez et al., 2004; Kiris et al., 2015).

These data suggests that BonT/A effects could be counteracted clinically by Src targeting drugs. This is important as serious or even lethal botulinic intoxications, while rare, still occur in developed countries like France (“Cas de botulisme alimentaire à Bordeaux : 15 cas recensés, dont 10 hospitalisés et 1 décès. Point de situation au 14 septembre 2023,” n.d.).

Another four genes are linked to the retromer complex, which controls endosomes to Golgi traffic. For BoNT/A, traffic from endocytic vesicle to Golgi requires retro-axonal transport, which is supported by the requirement for dynein. Previous studies had shown retro-axonal transport for BoNT/A (Harper et al., 2016; Restani et al., 2012). By extension, these results suggest that SV2 is also transported retro-axonally (Lund et al., 2021; Tanner et al., 1996). Consistently, we found that SV2 accumulates at neurite tips when the retromer complex is silenced. SV2 is implicated in epilepsy and various neurodegenerative diseases such as Alzheimer’s and Parkinson’s (Ciruelas et al., 2019; Stout et al., 2019).

Another complex required is the translocon. When Sec61G is depleted, mNG-complementing BoNT/A still accumulates in the ER but is unable to reach the cytosol, indicating that the Sec61 complex is involved in retrotranslocation. While the best-described function is the co-translational insertion of proteins in the ER, Sec61 has also been implicated in retrotranslocation linked to ER-associated degradation (ERAD) (Römisch, 2017). Sec61 thus likely functions as a bidirectional channel for proteins (Römisch, 2017).

Overall, we propose that the BoNT/A-SV2 complex is internalised by ReNcell VM cells at the neurite tip where SV2 is exposed. While we did not demonstrate this point formally in RenCell VM cells, it is well established in other neurons and it is consistent with the increased toxin sensitivity after neuronal stimulation **(Supplementary Figure 5A)**. After internalisation, instead of being translocated from endocytic vesicles, the toxin is retrogradely trafficked to the soma-located Golgi in a retromer-dependent fashion **(Supplementary Figure 5B).** After trafficking through the Golgi, the toxin is transported to the ER **(Supplementary Figure 5C)**. There, BoNT/A LC translocates via the Sec61 translocon into the cytosol. It is likely that LC detachment from ER membranes requires the separation of heavy and light chains through the activity of thioredoxin reductases and chaperones **(Supplementary Figure 5C)**. Thus, in neurons in culture, BoNT/A LC likely cleaves SNAP25 first in the soma before reaching the neurite terminals where it blocks fusion of synaptic vesicles at the plasma membrane **(Supplementary Figure 5D).** The location of the Golgi apparatus virtually exclusively in the soma of differentiated Ren-VM cells explains why the toxin has to traffic to the soma. This trafficking in turn explains the 48h between addition of the toxin and full cleavage of SNAPR.

Our study contradicts the long-established model of BoNT intoxication, which is described in several reviews specifically dedicated to the subject (Dong et al., 2019; Pirazzini et al., 2017, 2016; Rossetto et al., 2021). In short, these reviews support the notion that BoNT are molecular machines able to mediate their own translocation across membranes. This notion has convinced some cell biologists interested in toxins and retrograde membrane traffic, who follows this model of BoNT mode of translocation in their reviews (Mesquita et al., 2020; Williams and Tsai, 2016).

But is this notion well supported by data? A careful examination of the primary literature reveals that early studies indeed report that BonTs form ion channels at low pH values (Donovan and Middlebrook, 1986; Hoch et al., 1985). These studies have been extended by the use of patch-clamp (Fischer et al., 2009; Fischer and Montal, 2007a). These works and others lead to various suppositions on how the toxin forms a channel and translocate the LC (Fischer and Montal, 2007b; Pirazzini et al., 2016).

However, only a single study claims to reconstitute in vitro the translocation of BonT LC across membranes (Koriazova and Montal, 2003). In this paper, the authors report one key experiment using a system of artificial membranes separating two aqueous compartments. They load the toxin in the cis compartment and measure the protease activity in the trans compartment after incubation. However, when the experimental conditions described are actually converted in terms of molarity, it appears that the cis compartment was loaded at 10^e-8^M BonT and that the reported translocated protease activity is equivalent to 10^e-17^ M (Figure 3D, (Koriazova and Montal, 2003)). Thus, in this experiment, about 1 LC molecule in 100 millions has crossed the membrane. Such extremely low transfert rate does not tally with the extreme efficiency of intoxication in vivo, even while taking into account the difference between artificial and biological membranes.

In sum, a careful analysis of the primary literature indicate that while there is ample evidence that BoNTs have the ability to affect membranes and possibly create ion channels, there is actually no credible evidence that these channels can mediate the translocation of the LC. As mentioned earlier, it is unclear how such a self-translocation mechanism would function thermodynamically. It is worth noting that a similar self-translocation model was proposed for other protein toxins such as Pseudomonas exotoxin, which have similar molecular organisation as BonT (Taupiac et al., 1999). However, it has since been demonstrated that the PE toxins require cellular machinery, in particular in the ER, for intoxication (Bassik et al., 2013; Moreau et al., 2011; Schäuble et al., 2014).

By contrast, our model proposes a mechanism without a thermodynamic problem, is consistent with current knowledge about other protein toxins, such as PE, Shiga and Ricin, and can help explain previously puzzling features of BonT effects.

## Acknowledgements and conflict of interest

This study was funded by a grant from IPSEN and by an Industry Alignment Fund Grant from A*STAR. Jacquie Maignel, Laurent Pons, and Matthew Beard are employees of Ipsen. Keith Foster and Omar Loss are former employees of Ipsen. F. Bard is currently funded by a grant “Leader in Oncology” from the Fondation “ARC pour la recherche sur le cancer” and by a Chaire d’Excellence from AMIDEX: AMX-20-CE-03.

## Methods

### Chemical reagents

The reagents used in this study are summarized in **Supplementary Figure 6A.** BoNT/A was synthesized as previously described (Stancombe et al., 2012).

### Constructs

Constructs used in this study are depicted in **Supplementary Figure 6B** were generated using gene synthesis and cloned into pDONR221 entry vector (GeneArt, ThermoFisher Scientific). The entry clones were subcloned into pLenti6.3-V5 destination vectors using gateway LR cloning (ThermoFisher Scientific). In the split-mNG fluorescence reconstitution system, the first 10 β-barrel helices of monomeric Neon Green (mNG) are fused with or without target proteins while the last beta-strand is fused with the protein of interest. Interaction of the two partners will reconstitute fluorescence (Luong et al., 2020). BoNT/A-mNG_11x3_ was synthesized by addition of 3 mNG_11_ tags flanked with GSGSG (Gly-Ser) linkers at the N-termini of BoNT/A. siGENOME Human siRNA libraries were purchased from Dharmacon (Horizon Discovery).

### Cell lines

ReNcell®VM human neural progenitor cell line (Sigma Aldrich) were maintained in ReNcell ®NSC Maintenance medium supplemented with 20ng/mL of epidermal growth factor, EGF and basic fibroblast growth factor, bFGF at 37°C with 5% CO_2_. Cells were differentiated in differentiation medium (maintenance medium without EGF and bFGF; and addition of 10ng/mL of both glial-derived nerve factor (GDNF) and brain-derived nerve factor (BDNF) for two weeks with a change of media every 3 days. Cells were seeded on culture flasks or plates precoated overnight at 4°C with 20ug/mL laminin in DPBS. The cells were subcultured approximately every five days (90% confluence) by detaching them with Accutase.

### Stable cell line generation

Stable cell lines were generated using ViraPower Lentiviral Packaging Mix together with the following lentiviral constructs containing SNAP25 were cotransfected into HEK293FT cells using Lipofectamine 3000 reagent. Following incubation of cells, supernatant containing lentivirus was harvested and cellular debris was removed by centrifugation. The virus was transduced in ReNcell® VM with polybrene reagent (8mg/ml) and removed after 24 hours. Fresh complete growth medium was added and transduced cells were sorted using fluorescence-activated cell sorting using tGFP signal and selected using 10ug/mL blasticidin in maintenance medium.

### High-throughput siRNA and BoNT/A intoxication screen assay

The siRNA library was spotted in 384-well black μClear plates (PerkinElmer). siRNAs were transiently transfected at a final concentration of 25 nM per well using 0.25 μl Lipofectamine RNAiMAX reagent in 7.25 μl of OptiMEM medium, according to the manufacturer’s protocol. siNT3 (Non-targeting 3) was chosen as a control siRNA due to its least toxic properties in ReNcell VM. After 20 min of complex formation, complexes were dispensed to the differentiated cells per well using a Multidrop Combi dispenser (Thermo-Fisher Scientific). Medium was removed at 72 h post-transfection using Integra VIAFLO 384 (Integra Biosciences AG, Switzerland), and cells were washed with phosphate-buffered saline (PBS) twice before incubated with 6nM BoNT/A in differentiation medium with 58 mM KCl and 2.2 mM CaCl2 for 48 h. siRNA screening was performed in duplicate.

### Immunofluoresence

After 48 hours of BoNT/A intoxication, cells were fixed with 4% paraformaldehyde and 2% sucrose in PBS for 30 minutes and permeabilized with 0.2% Triton X-100 for a further 10 minutes. The cells were then stained with primary antibody diluted in 2%FBS in PBS for 2 hours. Cells were subsequently washed three times for 5 minutes with 2%FBS in PBS and stained for 1 hour with secondary antibody conjugated with fluorophore and Hoechst 33342 diluted in 2% FBS in PBS. The cells were then washed three times for 5 minutes with PBS before imaging.

### Imaging and image analysis

High-throughput imaging was carried out using an automation-enabled Opera Phenix system with 20X objective (PerkinElmer). GFP, RFP and Hoechst channels were imaged and image analysis was done using Harmony and Columbus software (PerkinElmer). In brief, nuclei counts were generated using the Hoechst channel while cell masks were obtained using the RFP channel. The cell-masked GFP channel was measured and GFP/RFP ratio readout was obtained.

### Data formatting and normalization

Genome-wide RNAi screen data was imported and analyzed in ScreenSifter software as previously described (Kumar et al., 2013). GFP/RFP ratio was normalized to controls and Z-score graphs were plotted.

### Western blot analysis

Rencell VM cells were transfected with siRNAs in a 10 cm dish for 3 days. On the third day, Cells were washed twice using ice-cold PBS before scraping in PBS. Cells were centrifuged at 300g for 5 min at 4°C and were lysed with ice-cold lysis buffer (50 mM Tris [pH 8.0, 4°C], 200 mM NaCl, 0.5% NP-40 alternative, 1mM DTT, and Complete Protease Inhibitor (Roche) for 30 min with gradual agitation before clarification of samples by centrifugation at 10,000 g for 10 min at 4°C. Samples were diluted in lysis buffer with 2X SDS loading buffer and boiled at 95°C for 2 min. They were then resolved by SDS-PAGE electrophoresis using bis-tris NuPage gels as per manufacturer’s instructions (Invitrogen) and transferred to PVDF membranes which was blocked using 3% BSA dissolved in TBST (50 mM Tris [pH8.0, 4°C], 150 mM NaCl, and 0.1% Tween 20) at room temperature for 1 hour before incubation with antibodies as manufacturer’s instructions.

## Supplementary Figure Legends

**Supplementary Figure 1. Differentiation and siRNA depletion dynamics in ReNcell VM. (A)** ReNcell VM differentiated for 2 weeks with staining for nuclei (in blue) and neuronal markers b3-tubulin MAP2 (green) and for oligodendrocyte marker CNPase (2’,3’-Cyclic-Nucleotide3’-Phosphodiesterase). Scale bars 20 μm. **(B)** ReD SNAPR cell line differentiated for 2 weeks with staining of b3-tubulin (white). Expression of SNAPR construct is detected in the RFP and GFP channels. Scale bars 50 μm. **(C)** Undifferentiated and differentiated ReNcell VM transfected with siNT3 (Non Targeting 3) siRNA or siRNA against thioredoxin reductase 1 (TXNRD1) for 3 and 5 days showing persistent TXNRD1 protein depletion. Scale bars 50 μm. **(D)** Undifferentiated and differentiated ReD SNAPR cells transfected with siNT3 or SNAP25 siRNA for 3 and 5 days showing reduction of the SNAPR reporter. Scale bars 20 μm.

**Supplementary Figure 2. Genome-wide RNAi screen data normalization and refinement. (A)** Genome-wide siRNA library plate replicates (Exp1 and Exp2) from Figure 2C **(B)** Individual replicates from Figure 2D. **(C)** Averaged nuclei count for each siRNA, 289 genes result in cell number below the cut-off of 4000 nuclei per well. Gene targets of siRNA most affecting cell survival are annotated in the graph. **(D)** The 289 genes from C are over-imposed on the SNAPR reporter results, 80 genes fall within the positive regulators and 8 genes with the negative regulators. All 88 genes were excluded from further analysis **(E)** Targeted siRNA deconvolution screen with positive and negative hits and siRNA control either non-toxin treated (NT3-) or toxin-treated (NT3+). Positive controls (TXNRD1+) replicates. The GFP/RFP threshold was set at 1.9 for positive regulators and 1.0 for negative regulators. Green data points are individual siRNAs that do not pass the threshold and do not confirm the pool result. Averaged GFP/RFP ratio for genes below or above threshold lines for positive and negative regulators that were removed from the hit lists. **(F)** RNA sequencing data from ReD SNAPR cells represented as log_2_CPM (counts per million) with an exclusion threshold set at CPM = 0.25 (log_2_CPM = −2, genes below are deemed as unexpressed).

**Supplementary Figure 3. Gene ontological analysis of positive hits from genome-wide intoxication screen. (A)** Cellular component **(B)** Molecular function **(C)** Biological Processes

**Supplementary Figure 4. Retromer is required for retro-axonal trafficking of the BoNT/A receptor. (A)** Cytosolic mNG_1-10_ expressing RenVM cells were incubated with 50 nM of BoNT/A-mNG_11_ and fixed at 0, 24, 36 and 48 hour timepoints. The MFI was quantified in 20 cells for each condition and time point, in the soma and neurites. Scale bar 20 μm. **(B)** ReNcell VM stably expressing Dendra-SV2 treated with siVPS35 for 3 days and imaged after fixation. Insets show axons of interest from each condition and arrowheads indicate enlarged SV2 puncta. Scale bar 20 μm and inset shown at 10 μm. **(C)** Method for quantifying number and size of SV2 puncta. tGFP image is background-subtracted, thresholded then analyzed for number and size of particles. Quantification of the number of SV2 puncta along every 50 μm segment of the axon starting from the axon tip from **B.** Quantification of area of SV2 puncta in whole axons from **B. (D)** Comparison of mean fluorescence intensities (MFI) at axons and axon tips from control and siVPS35-treated cells. Quantification of axon tip/axon MFIs. Graphs are obtained from 30 axons across 3 independent experiments. Scale bar 10 μm.

**Supplementary Figure 5. Schematic model for intracellular trafficking of BoNT/A. (A)** BoNT/A HC binds to its cognate receptor SV2. Surface expression of SV2 is regulated by a cohort of enhancers and repressors. **(B)** BoNT/A-containing endosomes are retro-axonally trafficked to the soma through the retrograde function of the retromer complex. **(C)** BoNT/A eventually traffics to the translocation-competent ER via the Golgi. **(D)** The ER SEC61 translocon complex facilitates LC translocation of BoNT/A from the ER lumen into the cytosol where the TXNRD1 and HSP complexes release and refold BoNT/A LC. The LC diffuses and cleaves SNAP25 in the soma. **(E)** Progressive diffusion of LC into the axons and axon terminals ultimately cleaves SNAP25 on neurotransmitter-containing vesicles, blocking neurotransmission at the synapse.

## Notes

### Competing Interest Statement

The authors have declared no competing interest.

### Summary of Updates

Correct mistakes in figures and figure legends; alterations in the Introduction and Discussion sections.

