## Supplementary Figures for "Botulinum toxin intoxication requires retrograde transport and membrane translocation at the ER in RenVM neurons"

Fig. S1

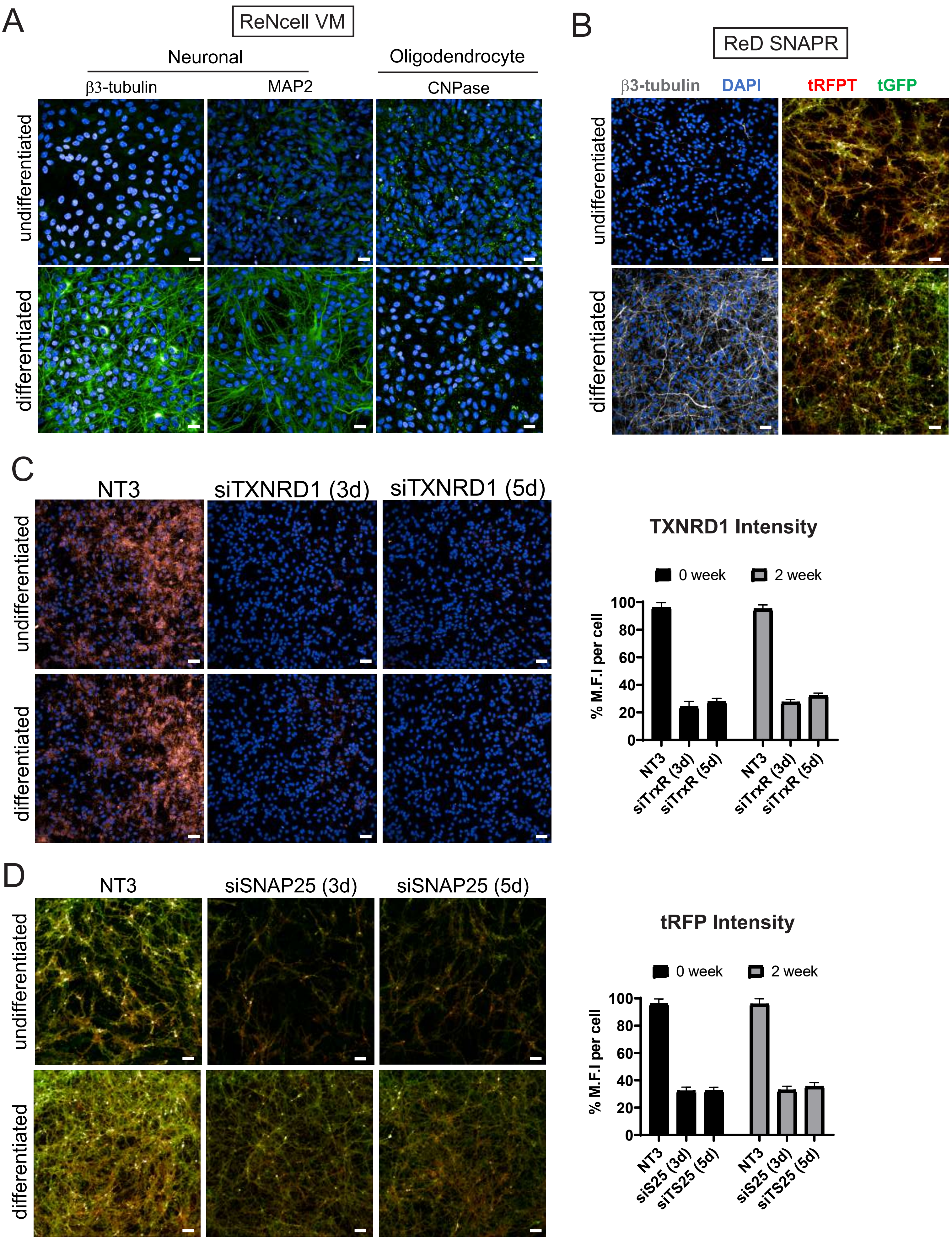
A
ReNcell VM
Neuronal
Oligodendrocyte
β3-tubulin
MAP2
CNPase
undifferentiated
differentiated
B
ReD SNAPR
β3-tubulin
DAPI
tRFPT
tGFP
undifferentiated
differentiated
C
NT3
siTXNRD1 (3d)
siTXNRD1 (5d)
undifferentiated
differentiated
TXNRD1 Intensity
0 week
2 week
% M.F.I per cell
100
80
60
40
20
0
NT3
siTrxR (3d)
siTrxR (5d)
NT3
siTrxR (3d)
siTrxR (5d)
D
NT3
siSNAP25 (3d)
siSNAP25 (5d)
undifferentiated
differentiated
tRFP Intensity
0 week
2 week
% M.F.I per cell
100
80
60
40
20
0
NT3
siS25 (3d)
siTS25 (5d)
NT3
siS25 (3d)
siTS25 (5d)

Fig. S2

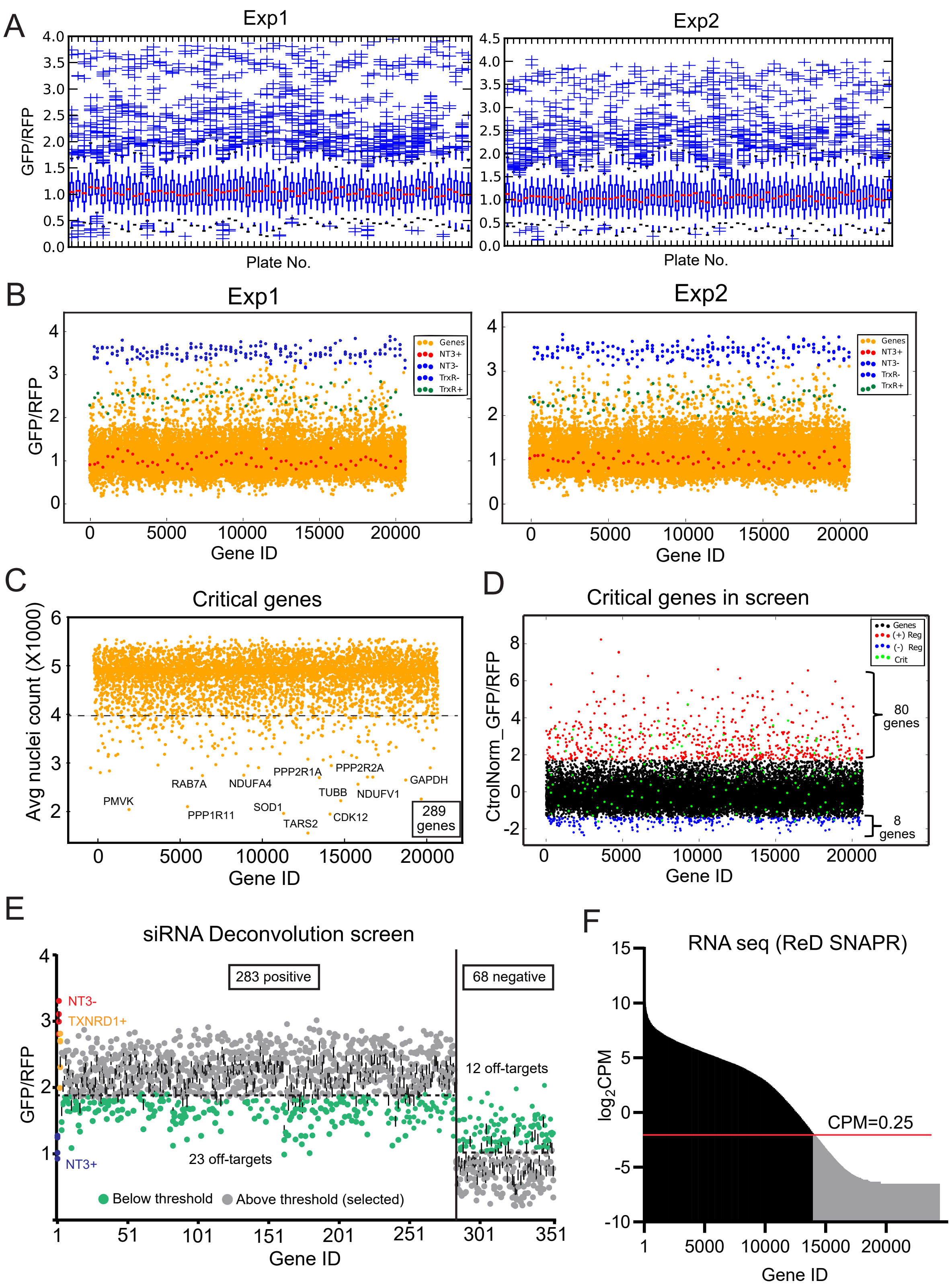
A
Exp1
Exp2
GFP/RFP
Plate No.
B
Exp1
Exp2
GFP/RFP
Genes
NT3+
NT3-
TrxR-
TrxR+
Gene ID
C
Critical genes
Avg nuclei count (X1000)
PMVK
RAB7A
NDUFA4
PPP2R1A
PPP2R2A
GAPDH
TUBB
NDUFV1
SOD1
PPP1R11
TARS2
CDK12
289 genes
Gene ID
D
Critical genes in screen
CtrolNorm_GFP/RFP
Genes
(+) Reg
(-) Reg
Crit
80 genes
8 genes
Gene ID
E
siRNA Deconvolution screen
283 positive
68 negative
NT3-
TXNRD1+
NT3+
12 off-targets
23 off-targets
Below threshold
Above threshold (selected)
GFP/RFP
Gene ID
F
RNA seq (ReD SNAPR)
log2CPM
CPM=0.25
Gene ID

Fig. S3

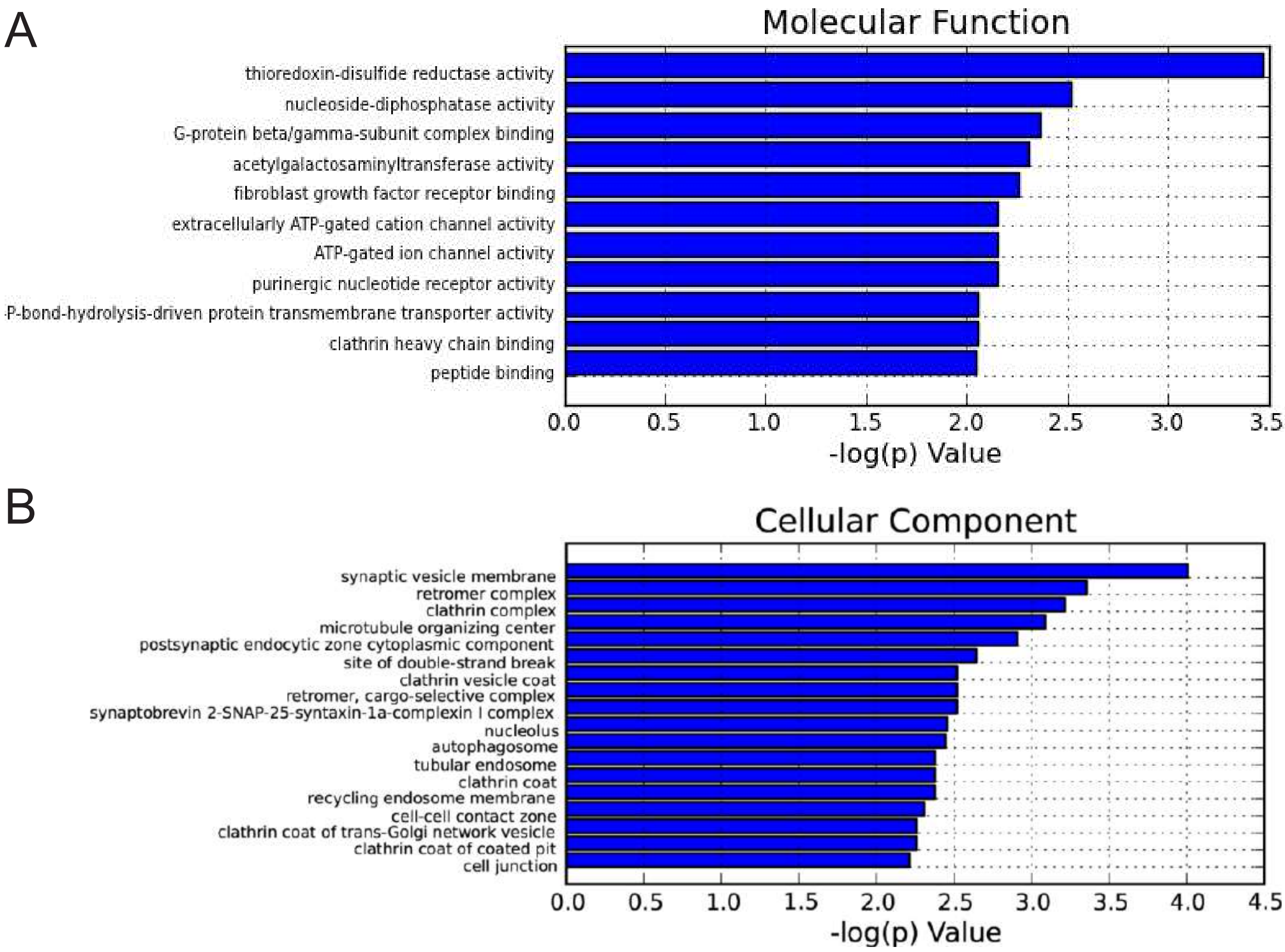
A
Molecular Function
thioredoxin-disulfide reductase activity
nucleoside-diphosphatase activity
G-protein beta/gamma-subunit complex binding
acetylgalactosaminyltransferase activity
fibroblast growth factor receptor binding
extracellularly ATP-gated cation channel activity
ATP-gated ion channel activity
purinergic nucleotide receptor activity
P-bond-hydrolysis-driven protein transmembrane transporter activity
clathrin heavy chain binding
peptide binding
0.0 0.5 1.0 1.5 2.0 2.5 3.0 3.5
-log(p) Value
B
Cellular Component
synaptic vesicle membrane
retromer complex
clathrin complex
microtubule organizing center
postsynaptic endocytic zone cytoplasmic component
site of double-strand break
clathrin vesicle coat
retromer, cargo-selective complex
synaptobrevin 2-SNAP-25-syntaxin-1a-complexin I complex
nucleolus
autophagosome
tubular endosome
clathrin coat
recycling endosome membrane
cell-cell contact zone
clathrin coat of trans-Golgi network vesicle
clathrin coat of coated pit
cell junction
0.0 0.5 1.0 1.5 2.0 2.5 3.0 3.5 4.0 4.5
-log(p) Value

Fig. S4

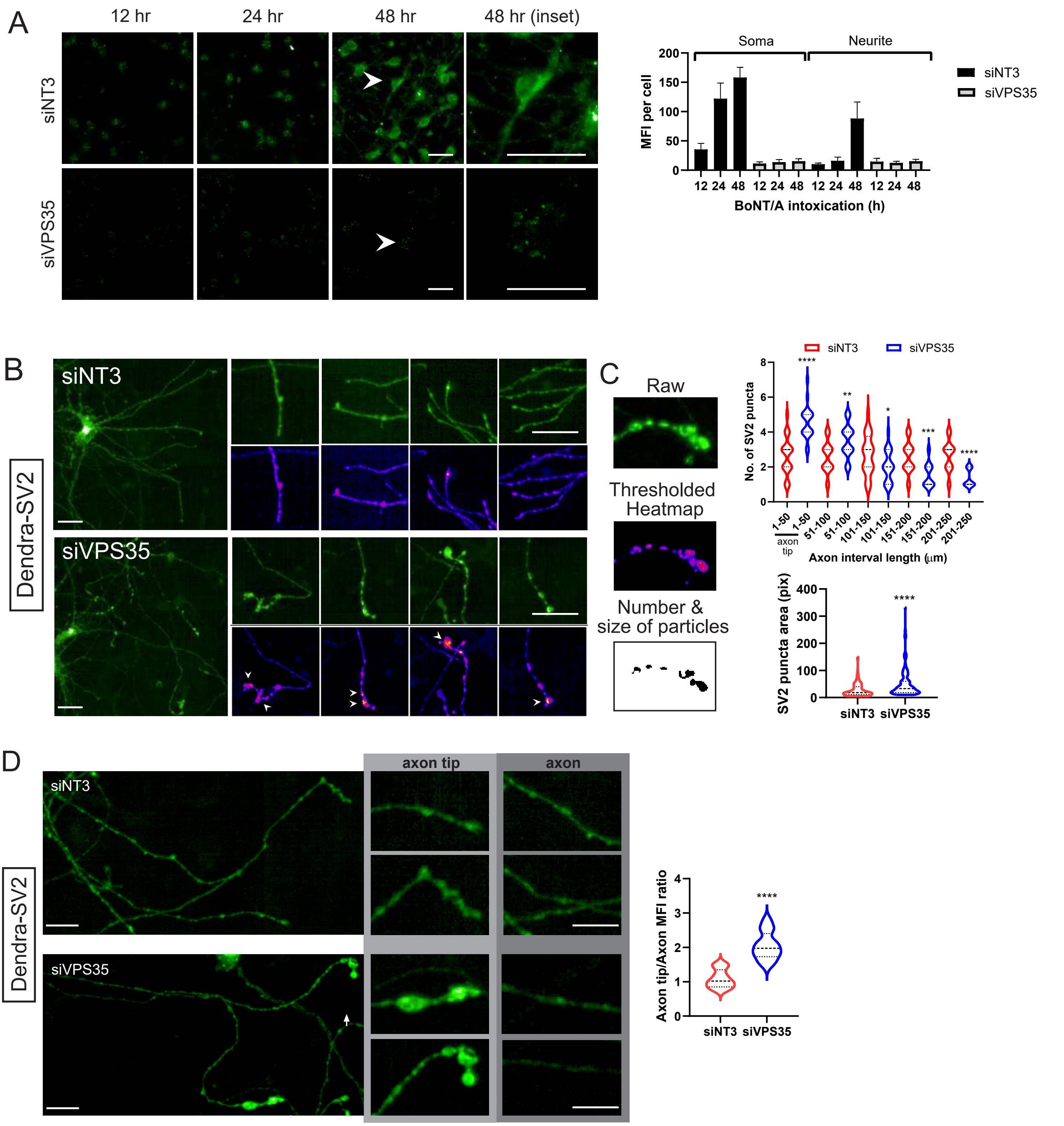
A
12 hr
24 hr
48 hr
48 hr (inset)
siNT3
siVPS35
Soma
Neurite
MFI per cell
BoNT/A intoxication (h)
siNT3
siVPS35
B
siNT3
siVPS35
Dendra-SV2
C
Raw
Thresholded Heatmap
Number & size of particles
siNT3
siVPS35
No. of SV2 puncta
Axon interval length (μm)
axon tip
SV2 puncta area (pix)
D
siNT3
siVPS35
axon tip
axon
Dendra-SV2
Axon tip/Axon MFI ratio

Fig. S5

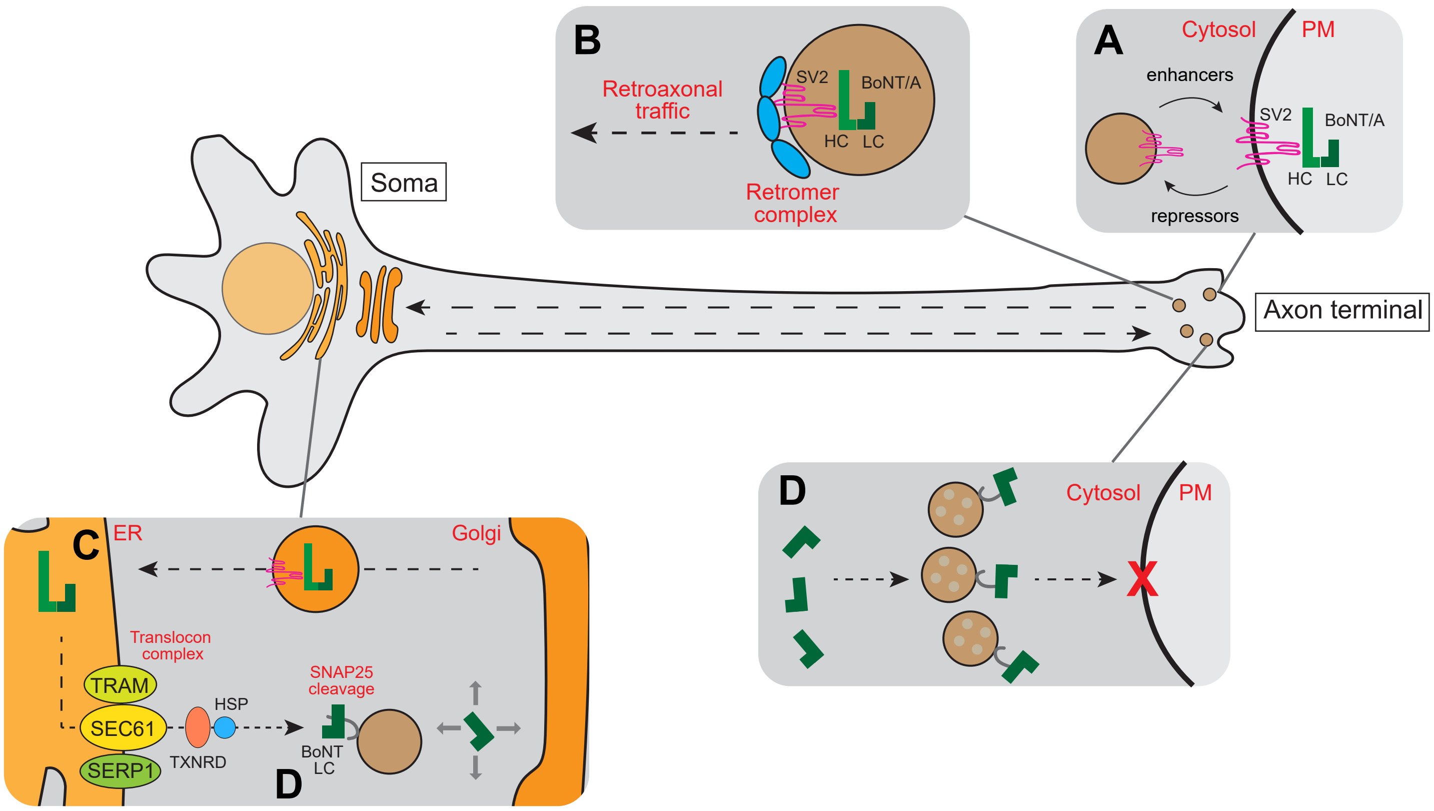
Soma
Axon terminal
B
Retroaxonal
traffic
SV2
BoNT/A
HC
LC
Retromer
complex
A
Cytosol
PM
enhancers
SV2
BoNT/A
HC
LC
repressors
C
ER
Golgi
Translocon
complex
TRAM
SEC61
SERP1
HSP
TXNRD
SNAP25
cleavage
BoNT
LC
D
D
Cytosol
PM

Fig. S6

A

| Reagent | Cat ID | Source |
|---|---|---|
| siGENOME pooled siRNA | | Dharmacon, USA |
| Deconvoluted siRNA | | Dharmacon, USA |
| Lipofectamine RNAimax | 13778150 | Invitrogen, USA |
| Lipofectamine 3000 | L3000015 | Invitrogen, USA |
| Human Epidermal Growth Factor | PHG0311 | Gibco, USA |
| Human Fibroblast Growth Factor-basic | PHG0261 | Gibco, USA |
| Human Nerve Growth Factor | A42578 | Invitrogen, USA |
| Human Brain-Derived Neurotrophic Factor | PHC7074 | Gibco, USA |
| ReNcell VM NSC Maintenance Medium | SCM005 | Sigma-Aldrich, USA |
| ReNcell NSC Freezing Medium | SCM007 | Sigma-Aldrich, USA |
| Laminin Mouse Protein | 23017015 | Gibco, USA |
| Accutase | A6964 | Sigma-Aldrich, USA |
| Potassium Chloride | P9541 | Sigma-Aldrich, USA |
| Calcium Chloride | C8106 | Sigma-Aldrich, USA |
| PBS, DPBS, Sterile water | | Gibco, USA |
| ViraPower Lentiviral Packaging Mix | A11146 | Invitrogen, USA |
| Polybrene | TR-1003 | Sigma-Aldrich, USA |
| Neuronal Marker IF Antibody Sampler Kit | 8752 | Cell Signaling, USA |
| SNAP25 antibody | 5308 | Cell Signaling, USA |
| tagRFPT antibody | AB233 | Evrogen, Russia |
| tagGFP antibody | AB121 | Evrogen, Russia |
| Thioredoxin Reductase 1 antibody | ab124954 | Abcam |
| SV2A antibody | 66724 | Cell Signaling, USA |
| SEC61G antibody | HPA053196 | Sigma-Aldrich, USA |
| Hoechst 33342 | H3570 | Invitrogen, USA |

B

**Constructs**

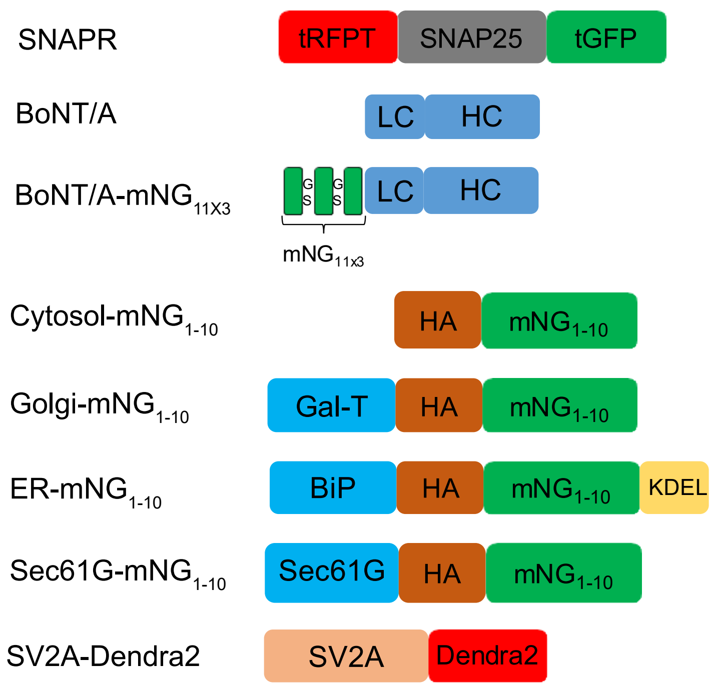
